# Mapping Early Brain Maturation: Anatomical Substrates of Cortical Connectivity Shifts in Neonates

**DOI:** 10.1101/2025.01.07.627373

**Authors:** Deying Li, Yufan Wang, Luqi Cheng, Simon B Eickhoff, Congying Chu, Lingzhong Fan

## Abstract

Understanding the early development of brain connectivity is essential for unraveling the mechanisms of brain maturation. In this study, we utilized diffusion MRI data from 242 term neonates to chart the developmental trajectory of cortical connectivity in infants. We quantified interareal variations in connectivity and identified three distinct global connectopies (GCs) along the rostrocaudal, dorsoventral, and mediolateral axes by using gradient mapping techniques. A pivotal age-dependent shift between the dorsoventral and mediolateral GCs occurred at 40 postmenstrual weeks. This shift was diminished by virtual thalamic fiber lesions, underscoring the critical role of thalamic projections. A longitudinal analysis disclosed a delayed structural connectivity pattern development in preterm infants who failed to exhibit the anticipated shift. This research helps to clarify the topographic principles that shape infant brain organization and emphasizes the influence of thalamic connectivity on early brain development. Our findings elucidate the anatomical underpinnings of early brain development, providing critical insights into the formation of cortical hierarchies and highlighting the influence of thalamic connectivity on developmental trajectories. These findings are essential for understanding typical brain growth and the pathogenesis of neurodevelopmental disorders.

## Introduction

The infant brain undergoes profound changes in its cortical microstructure ^[1, 2]^, macroscale functional organization ^[3, 4]^, and gene expression ^[5, 6]^ as it begins interacting with the external environment. As the neonatal brain adapts to different sensory inputs, this shift from the intrauterine to extrauterine environment initiates rapid brain growth, including the proliferation of dendritic arborizations ^[7, 8]^, accelerated growth of axonal myelination ^[9, 10]^, and a remarkable increase in synaptic density ^[11, 12]^, particularly during the early postnatal weeks. These microscale processes lead to neuroanatomical changes in key white matter tracts, with distinct patterns of size and microstructural variations observed across different fiber pathways ^[13–15]^. Despite an initial overproduction of connections that are later refined through activity-dependent pruning ^[16]^, the gene-regulated white matter maintains a high degree of organization. This organization is characterized by the spatial coordination of cortical areas within the connected brain ^[17]^, thereby revealing the principles of structural connectivity topography that underlie infant brain organization. In addition to the influence of genetic regulation and molecular signaling ^[18, 19]^, previous studies have highlighted the critical role of thalamocortical projections in cortical development and arealization both prenatally and postnatally ^[20, 21]^. Acting as a central relay for sensory input, the thalamus transmits external signals that drive early synaptic plasticity and refine the functional layout of the neocortex ^[22]^, gradually shaping a "protomap" of cortical organization ^[23]^. Thalamocortical projections are also directed to specific cortical areas during development under the guidance of transcription factors and axon guidance cues, facilitating the regional differentiation of the neocortex ^[24]^. While increasing a evidence has documented transformative white matter changes during this period ^[13–15]^, the specific contribution of thalamocortical connectivity to interareal connectivity variation is yet to be explored. Moreover, given the varying severities of cerebral white matter injury that are observed in preterm infants around their term-equivalent ages and are manifested as disrupted myelination patterns ^[25, 26]^, it is crucial to investigate the potential impact of exposure to preterm conditions on the structural connectivity landscape of the developing brain.

To address these unresolved questions, we used a substantial diffusion MRI dataset from the Developing Human Connectome Project (dHCP), encompassing 242 term neonates aged 38 to 43 postmenstrual weeks (PMW) at scan time, to quantify variations in interareal connectivity. Leveraging a gradient approach^[17]^, we identified three global connectopies (GCs) running along the rostrocaudal, dorsoventral, and mediolateral axes of the brain. Our results revealed a transition between the dorsoventral and mediolateral axes around 40 PMW. Considering the postnatal importance of thalamic projection tracts, we employed a virtual lesion approach to assess their influence on infant GCs. The 40 PMW transition ceased after we applied a virtual lesion of the thalamic projection tracts. Furthermore, an longitudinal analysis of the dataset revealed the impact of preterm birth on brain structural connectivity, with preterm infants showing delayed development and not demonstrating the transition at the expected time. This study, therefore, helps to elucidate the topographic principles that shape infant brain organization and offers valuable insights into the development of structural connectivity. The logic of our analysis is outlined in Figure 1, which provides an overview of the study’s methodology.

**Figure 1.**
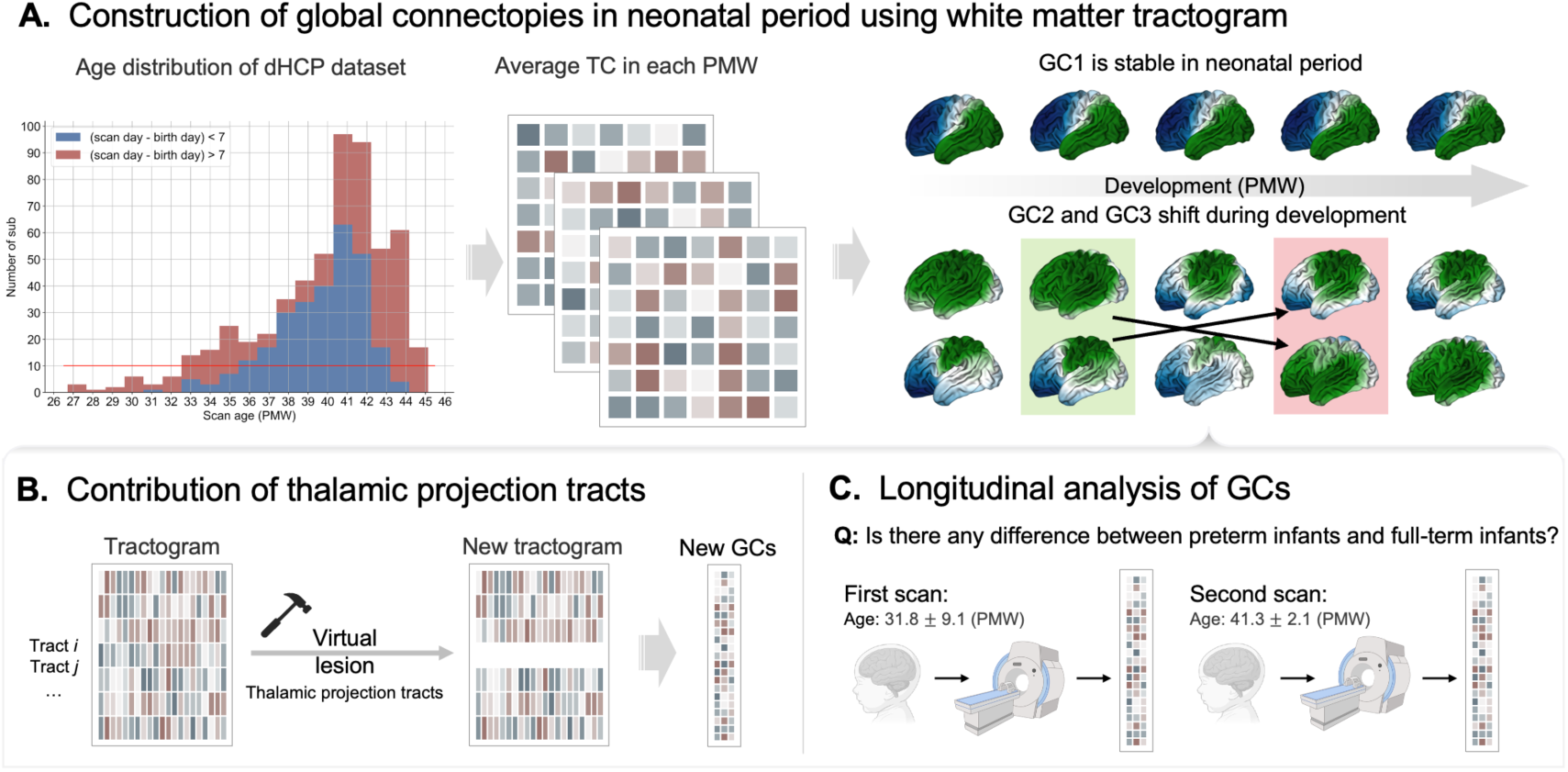
Overview of the analysis pipeline. **(A)** Global connectopies (GCs) in infants. **Left:** Postmenstrual age distribution of the dHCP infant cohort. **Middle:** White matter tractograms were used to calculate a similarity matrix called the tractogram covariance (TC) matrix. Diffusion embedding was performed on the TC matrix across different postmenstrual ages (from 38 to 43 postmenstrual weeks), resulting in low-dimensional gradients. **Right:** In infants, the first principal gradient follows a rostrocaudal pattern, while the second and third GCs exhibit dorsoventral and mediolateral patterns, respectively, with a notable shift occurring around 40 PMW. **(B)** Analysis of white matter bundles contributing to GCs. We conducted a virtual lesion analysis of the thalamic projection tracts to explore these contributions further. **(C)** Longitudinal analysis of GCs. Infants with two scans were included in this part. We calculated the group-level global connectopies for each of the two scans separately to investigate the impact of preterm birth on brain structural connectivity.

## Results

### Global connectopies in the human neonate brain

To investigate how the spatial organization of the human neonate brain is influenced by the underlying white matter structural connections and how it evolves with age postnatally, we characterized global connectivity across the entire brain for each postmenstrual week (PMW). Neonate data from the dHCP dataset ^[27]^ focused on individuals scanned between 38 and 43 PMW, as 37 PMW is the threshold for full-term delivery. We included only those individuals whose birth age and scan age differed by less than seven days (one week), as depicted in Figure 1A (Left, blue bar).

We constructed a vertex-wise similarity matrix of structural connectivity by correlating profiles from probabilistic tractography across approximately 30,000 vertices per hemisphere for each subject. We then computed tractogram covariance (TC) matrices, thresholded them at zero, and averaged them across subjects for each postmenstrual age. Thus, the TC matrix captured the structural connectivity similarity across the brain.

We applied diffusion embedding, a manifold learning method previously used to identify functional gradients ^[28]^, to the TC matrix. The resultant components revealed the positional relationships between the vertices along the axes with the most dominant structural connectivity differences. Although the order of global connectopies changed with development, the first three global connectopies exhibited rostrocaudal, mediolateral, and dorsoventral patterns (Figure 2). Specifically, the rostrocaudal axis (GC-RC) varied along a rostrocaudal axis, radiating from the occipitoparietal cortex and ending in the prefrontal cortex. The mediolateral axis (GC-ML) displayed a mediolateral pattern, with the highest expression in the lateral temporal and prefrontal cortex and the lowest in the cingulate cortex. The dorsoventral axis (GC-DV) followed a dorsoventral pattern, anchored at one end by the occipital, inferior temporal, and orbitofrontal cortex and at the other end by the sensorimotor cortex (Figure 2). Given the limited number of subjects in the early stages (fewer than 10, Figure 1A Left), we expanded the allowable postmenstrual age difference between birth and scan to 28 days (4 weeks) (Figure S1) and recalculated the global connectivities from 33 PMW to 43 PMW (with more than 10 subjects) (Figure S2). We found that the global connectivities remained stable from 38 PMW to 43 PMW.

**Figure 2.**
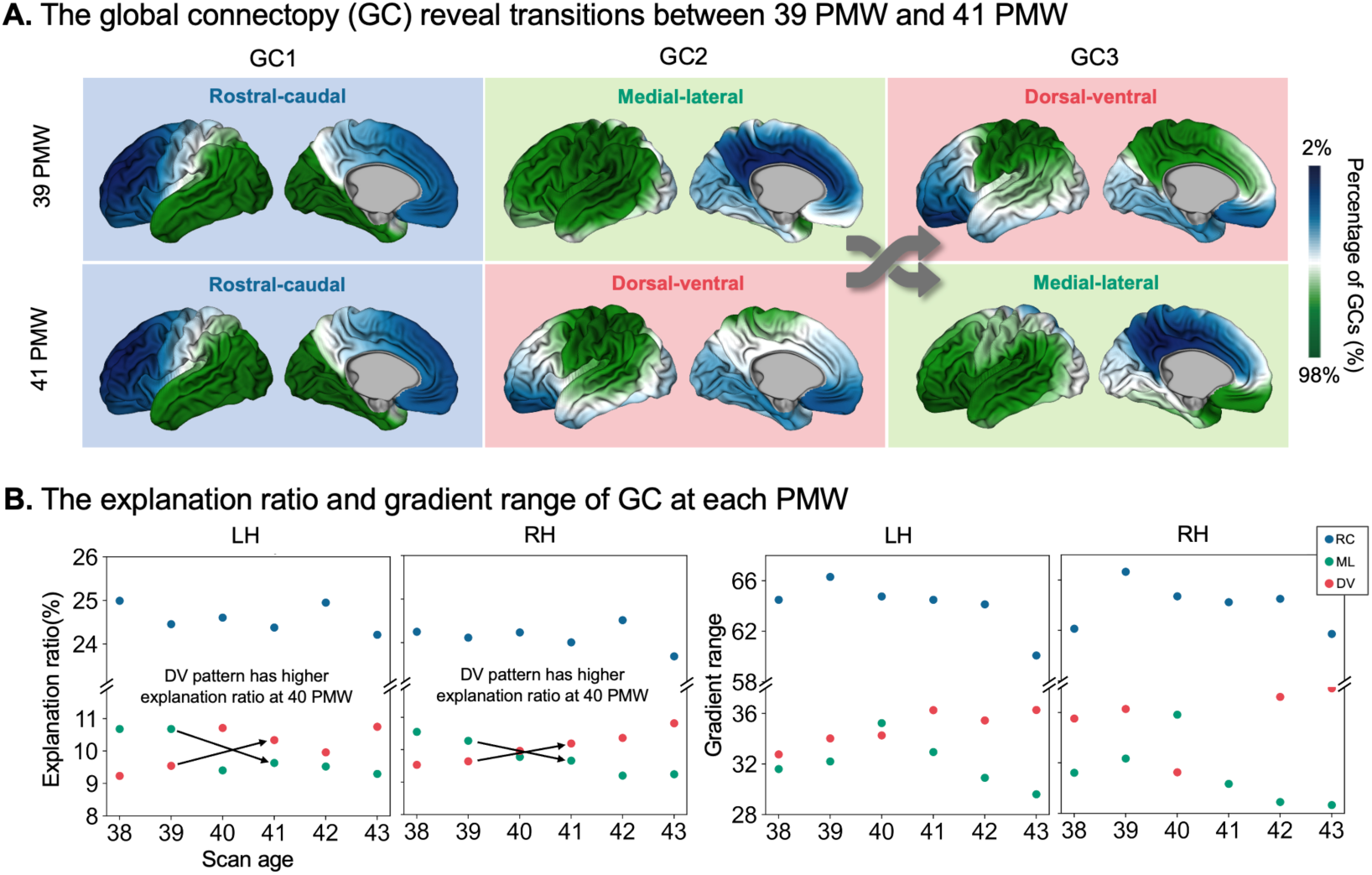
The three global connectopies after the birth of human infants. **(A)** The three global connectopies ran rostrocaudally, mediolaterally (green box), and dorsoventrally (red box) in 39 postmenstrual weeks (PMW) and 41 PMW. The dominant global connectopy maintained stable, while the second axis shifted from a mediolateral to a dorsoventral pattern at 41 PMW. **(B) Left:** The rostrocaudal pattern was the dominant global connectopy from 38 to 43 PMW, shifting from mediolateral to dorsoventral at 40 PMW. **Right:** The gradient range followed a trend that was similar to that of the explanation ratio with the development of the highest rostrocaudal pattern and a shift occurring at 40 PMW.

Our findings in infants contrasted with those in adults ^[17]^ in that the predominant global connectivity pattern in infants was rostrocaudal, whereas in adults, it was dorsoventral. Notably, the secondary axis in infants transitioned from mediolateral to dorsoventral around 40 PMW (Figure 2, Figure S2). These age-dependent shifts suggest a reorganization of structural connectivity during late neonatal development. Similar patterns were observed at the individual level for GCs. We discovered that GC1 was highly stable across individuals. Approximately 60% of infants exhibited stable GC2 and GC3 before 39 PMW. Moreover, over two-thirds of the infants underwent a transition in their GC patterns after 40 PMW, consistent with the group-level GC trajectories.

To evaluate the development of global connectopies across age, we calculated the explanation ratio (i.e., the variance accounted for by global connectopies) and the gradient range (i.e., the difference between the values of the positive and negative extremes of the gradient axis). The variation in the explanation ratio metric sheds light on one of the factors explaining this maturation jump. Unlike the rostrocaudal pattern, which consistently retained a high explanation ratio, the explanation ratio of the mediolateral pattern gradually decreased, while that of the dorsoventral pattern gradually increased. At 40 PMW, the dorsoventral pattern exceeded the mediolateral in the explanation ratio, emerging as the second most significant pattern in both hemispheres (Figure 2B Left). This transition was further validated by the gradient range curves across ages (Figure 2B Right). In conclusion, the rostrocaudal axis had already emerged and remained stable prenatally, while the other two axes exhibited a substantial increase around the late-term age, eventually stabilizing, as indicated by these two measures.

### Thalamic projection tracts support global connectopies shift

Considering the significant expansion of white matter during early development, we focused on identifying the specific tracts associated with the emergence of global connectopies. Previous studies have emphasized the crucial role of thalamic projection tracts in postnatal brain development ^[29, 30]^. Building on these findings, we examined the role of these tracts in our research. Specifically, we reconstructed thalamic projection fiber bundles using TractSeg ^[31]^, including ATR (anterior thalamic radiation), OR (optic radiation), STR (superior thalamic radiation), T PREF (thalamo-prefrontal), T PREM (thalamo-premotor), T PREC (thalamo-precentral), T POSTC (thalamo-postcentral), T PAR (thalamo-parietal), and T OCC (thalamo-occipital).

To evaluate the significance of these tracts, we conducted a virtual lesion analysis. We set the probability values for all the thalamic projection tracts to zero within the tractogram and recalculated the global connectivities using multi-modal parcellation ^[32]^ (virtual lesion GCs, vlGCs). This manipulation revealed that the observed shift was no longer present (Figure 3). Consequently, our findings highlight the pivotal role of thalamic projection tracts in early brain development, particularly in the formation of the dorsoventral pattern.

**Figure 3.**
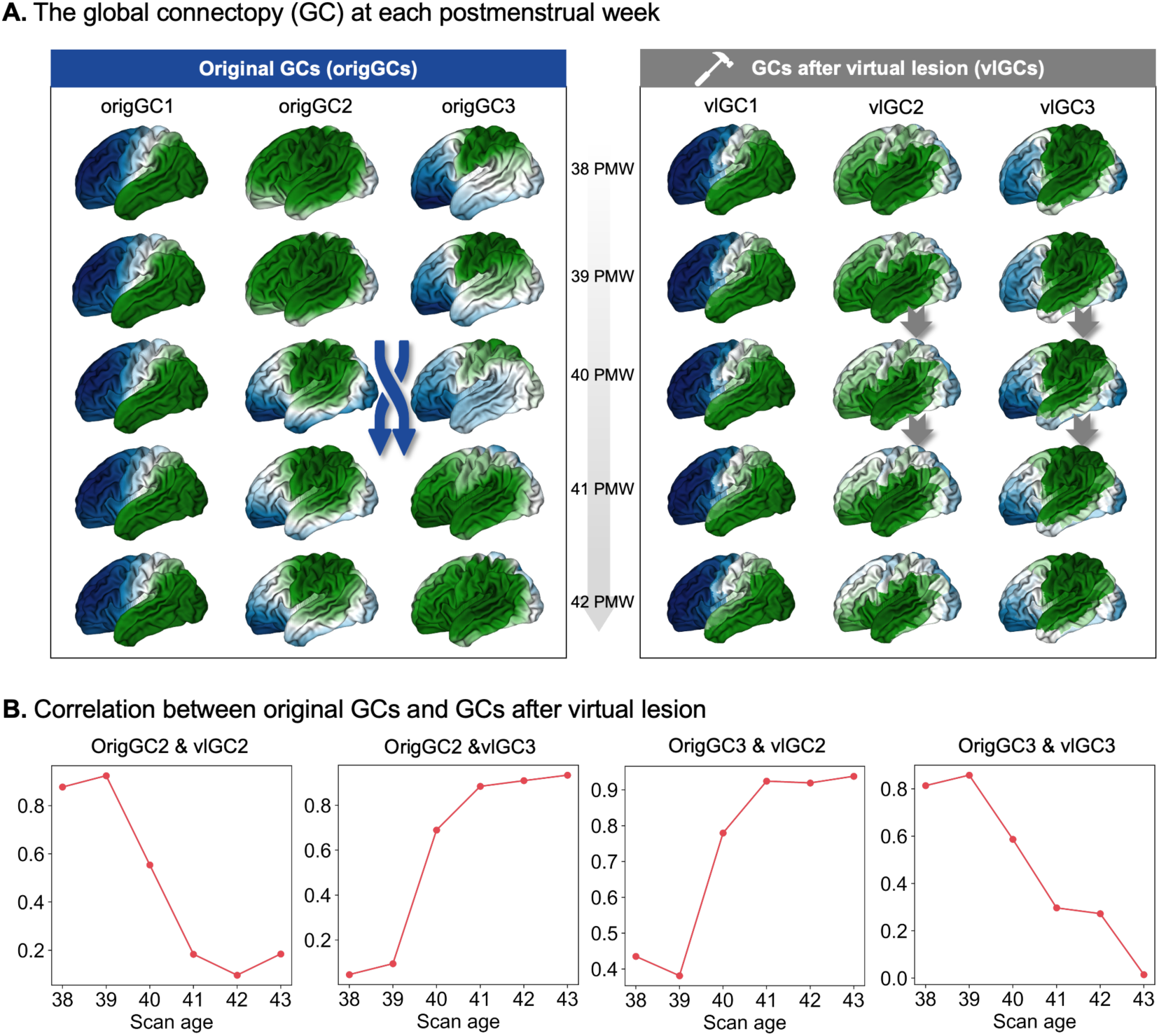
The global connectopies after a virtual lesion of thalamocortical tracts from 38 PMW to 43 PMW. **(A) Left:** The original global connecopies (OrigGCs) from 38 PMW to 42 PMW. From 39 PMW to 41 PMW, a transition can be observed in GC2 and GC3. **Right:** By setting the probability of all the thalamic projection tracts to zero within the tractogram, we effectively simulated the absence of these key white matter pathways. The resultant global connectivities (virtual lesion GCs, vlGCs) failed to demonstrate the typical transition between GC2 and GC3, indicating that these tracts are not only necessary for the formation of global connectopies but also for their dynamic shifts. **(B)** The correlation between the OrigGCs and the vlGCs after a virtual lesion reveals distinct patterns. OrigGC2 shows a high correlation with vlGC2 before 40 PMW, but after 40 PMW, it exhibits a high correlation with vlGC3. Similarly, OrigGC3 shows a high correlation with vlGC3 before 40 PMW, but after 40 PMW, it exhibits a high correlation with vlGC2.

### Impact of preterm birth on global connectopies

In our initial findings, we concentrated on full-term infants, ensuring that their birth age and scan age were within a week to exclude preterm infants. Recognizing the profound impact of preterm birth on infant development ^[25, 26]^, we extended our analysis to preterm infants who had undergone two scans to probe its effects on global connectopies further. Our longitudinal analysis encompassed 43 preterm infants with two scans. For the subsequent analysis, we selected 36 infants whose second scan was conducted after 40 postmenstrual weeks (PMW), as detailed in Figure 4A. The global connectopies were calculated using the methodologies described above.

**Figure 4.**
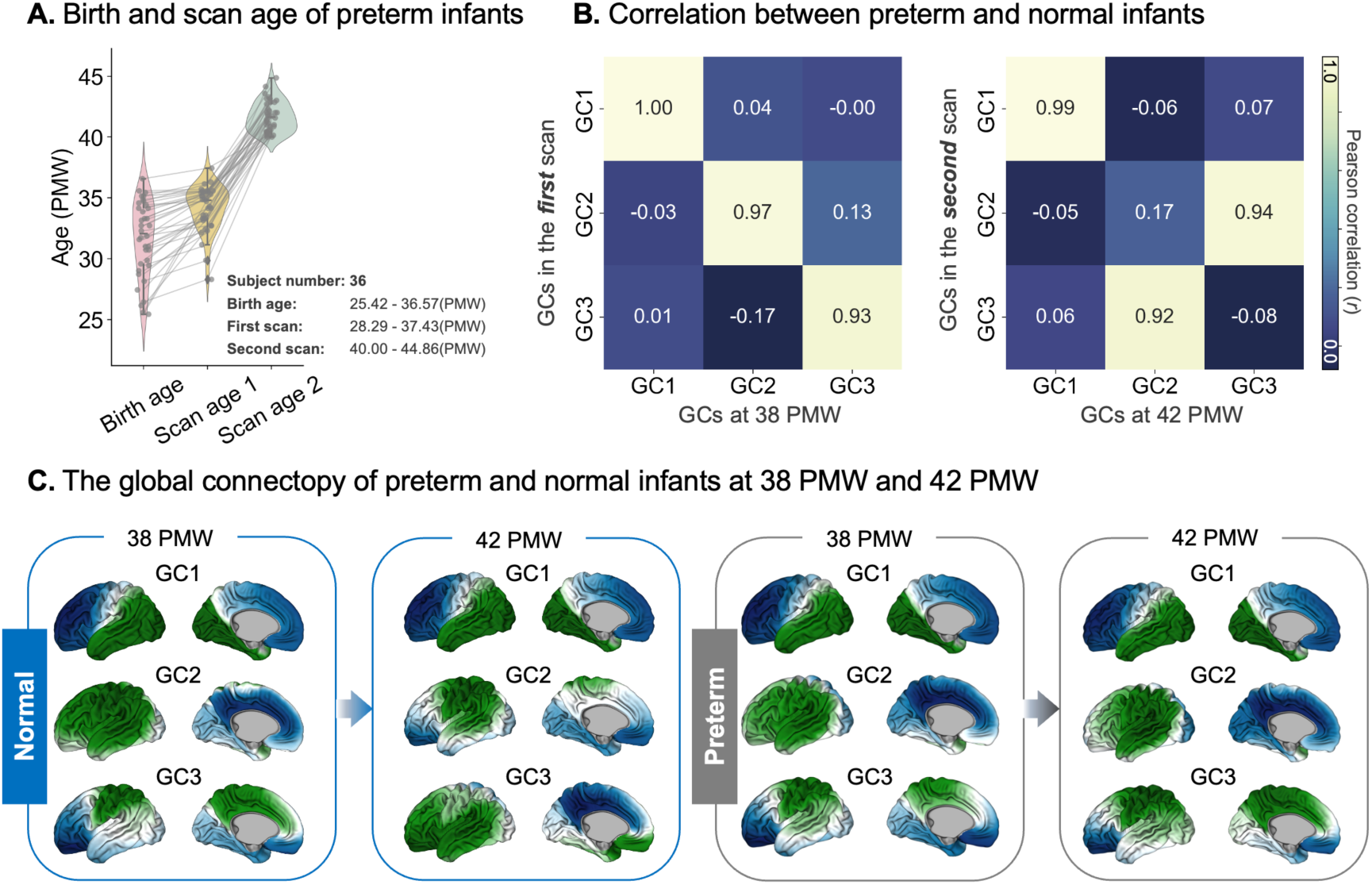
Longitudinal assessment of preterm birth impacts on global connectopies. **(A)** The analysis included 36 infants from the dHCP dataset who underwent two scans. The birth ages of these infants ranged from 25.42 to 36.57 PMW, with the first scan conducted between 28.29 to 37.43 PMW and the second scan between 40.00 to 44.86 PMW. **(B) Left panel:** We observed a strong correlation between the group-level global connectopies identified in the first scan and those at 38 PMW in full-term infants, with correlation coefficients of GC1: *r* = 1.00, GC2: *r* = 0.97, and GC3: *r* = 0.93. **Right panel:** a high correlation was found between the group-level global connectopies from the second scan and those at 42 PMW in full-term infants (GC1: *r* = 0.99; GC2 from the preterm group and GC3 from full-term group: *r* = 0.94; GC3 from the preterm group and GC2 full-term group: *r* = 0.92). **(C)** The global connectopies of preterm and normal subjects at 38 PMW and 42 PMW are compared. Normal subjects exhibit a transition from 38 PMW to 42 PMW, whereas preterm subjects show no such transition.

We observed that preterm infants showed similar global connectopic patterns to full-term infants before 39 PMW, but their development decelerated post-40 PMW. Given the stability of global connectopies from 33 PMW to 38 PMW (Figure S2), we calculated the correlation between the group-level global connectopies from the first scan and those at 38 PMW in full-term infants, revealing strong correlations (Figure 4B Left; GC1: *r* = 1.00; GC2: *r* = 0.97; GC3: *r* = 0.93). However, the global connectopies from the second scan of preterm infants displayed a misalignment with those at 42 PMW. Specifically, GC2 from preterm infants correlated highly with GC3 from full-term infants (r = 0.94), and GC3 from preterm infants correlated highly with GC2 from full-term infants (r = 0.92; Figure 4B Right).

## Discussion

This study elucidates the intricate developmental sequence of global connectopies in neonates, a critical phase in early brain maturation. To bridge knowledge gaps, we systematically mapped the cerebral connectivity in full-term neonates, revealing the emergence of adult-like rostrocaudal, mediolateral, and dorsoventral patterns ^[17]^. Notably, a transition in connectivity dominance around 40 postmenstrual weeks indicates a significant reorganization in cortical organization. Thalamic projection tracts, crucial for postnatal maturation, were found to influence these connectopies, especially the dorsoventral axis, substantially. Preterm infants showed delayed connectivity stabilization, emphasizing the impact of preterm birth on brain development. Our findings underscore the anatomical underpinnings of early brain development, offering insights into cortical hierarchy formation and the potential origins of neurodevelopmental disorders.

The global connectopies derived from white matter partially characterize the low-dimensional embedding of the neonatal cortex’s connectivity architecture, which is organized along the same axes as in those adults ^[17]^. This alignment of cortical connectivity patterns with the primary genetic and spatiomolecular axes of brain development ^[33, 34]^ and the chemotactic gradients established during early embryogenesis ^[35, 36]^ suggests that there are conserved principles governing cortical rewiring as the cortex develops. Axonal growth, a complex process integral to forming association projections between cortical areas, is influenced by various factors, including chemotactic gradients and molecular cues that direct the growth and projection of axons ^[37, 38]^. Moreover, radial glial cells in the embryonic brain, which facilitate the generation, placement, and allocation of neurons in the cortex and regulate how they wire up, may act as "highways" for axons, scaffolding their growth and ensuring their proper trajectories ^[39, 40]^. These observations indicate that the genetic blueprint present early in brain development is instrumental in guiding the formation of neural circuits and the arealization of the cortex ^[17, 41, 42]^.

Interestingly, the neonatal cortex predominantly exhibits a rostrocaudal pattern, contrasting with the dorsoventral pattern typically observed in adults. This distinction can be attributed to several developmental factors. A key contributing factor may be the extended neurogenic period in humans, which is longer than in species with smaller brains, resulting in a preferential expansion of structures that develop later ^[43–46]^. Supporting this, evidence indicates that cortical neurogenesis progressively terminates along the rostrocaudal axis ^[47–49]^. Consequently, the earlier cessation of neuronal production in the anterior part of the cerebral cortex may foster conditions conducive to increased neuronal size and dendritic arborization ^[46, 50, 51]^. Despite the limitations of our data, which starts around 30 PMW, the stable dominance of the rostrocaudal pattern during this period implies that it likely emerged even earlier. Indeed, thalamocortical tracts, which significantly contribute to this pattern, have been identified as early as 23 PMW ^[52]^, demonstrating an anterior-posterior gradient in cortical connectivity. This gradient is consistent with the topographical organization of thalamic nuclei observed in rodent histological studies ^[24, 53]^. Furthermore, the variations in neurogenesis along the rostrocaudal axis are mirrored in the decreasing neuron density and increasing neuronal connectivity from the posterior to the anterior brain areas ^[54]^, potentially accounting for the anterior-posterior functional gradient observed in the late neonatal period ^[55]^.

The shift from the dorsoventral to mediolateral axes around 40 postmenstrual weeks indicates a critical phase of gradual developmental changes in the neonatal brain, underpinned by the reorganization of cortical circuitry. Toward the end of gestation and throughout the first two postnatal months, dendrites, spines, and synapses are rapidly developed, particularly in cortical layer IV, which is the primary recipient of thalamocortical projections ^[56–58]^. The maturation of the thalamocortical circuitry and its interaction with sensory inputs are instrumental in sculpting the cortical topological organization ^[59, 60]^. While not the focus of this study, the ongoing development of these connections during childhood and adolescence is essential for functional specialization ^[21, 61]^, potentially influencing the shift in dominant global connectopies observed in adults. The increased significance of the dorsoventral pattern likely mirrors the processes of neural pruning and synaptic strengthening, which prepare the brain to engage with the abundant sensory and motor stimuli encountered prenatally and postnatally ^[55]^. These developmental mechanisms are crucial for refining neural circuits for efficient information processing and integrating new experiences during this sensitive period. Our findings show that the dorsoventral pattern’s significance did not escalate postnatally when thalamic projection tracts were virtually lesioned (Figure 3), underscoring the pivotal role of these tracts and, by extension, external stimuli in the formation of the neonatal brain’s white matter organization. Additionally, our longitudinal analysis revealed developmental delays associated with preterm birth, possibly due to impairments in thalamic projection tracts ^[25, 26]^.

Finally, the current work must acknowledge several technical and methodological limitations. The primary concern revolves around the precision of structural connectivity mapping utilizing diffusion tensor imaging ^[62]^. Despite this, diffusion tractography is an indispensable technique for the in vivo, non-invasive characterization of white matter throughout the brain, particularly in fetal and neonatal contexts. The challenges inherent in this approach can be partially addressed through integrated microscopy data analysis, enhancing the accuracy of the findings ^[63, 64]^. While the distance effect is a factor to consider when mapping structural connectivity patterns, existing evidence indicating a correlation between global connectivities and genetic topography suggests that these patterns may transcend distance dependencies ^[17]^. This insight implies that the observed connective patterns are not solely a function of the spatial relationships between brain regions. Furthermore, the interplay between connectivities and brain geometry merits closer scrutiny to elucidate the interrelationship between developing brain tissues that are changing at different rates. Extending the analysis of global connectopies to encompass earlier prenatal stages and through childhood and adolescence could yield invaluable insights into the dynamics of cortical organization during critical development periods.

## Methods

### Data and preprocessing

#### Data collection

A total of 886 datasets (783 neonatal subjects) from the third release of the Developing Human Connectome Project (dHCP) database were used (https://www.developingconnectome.org/) ^[27]^. The dHCP was approved by the UK Health Research Authority. Informed parental consent was obtained for imaging acquisition and data release.

The scanning procedures and acquisition parameters were detailed in previous publications ^[65]^. In brief, all infants were scanned without sedation in a scanner environment optimized for neonatal imaging, including a dedicated 32-channel neonatal coil on a 3T Philips Achieva. MR-compatible ear putty and earmuffs were used to provide additional noise attenuation. Infants were fed, swaddled, and positioned in a vacuum jacket before scanning to promote natural sleep. All scans were supervised by a neonatal nurse and/or pediatrician who monitored heart rate, oxygen saturation, and temperature throughout the scan.

T2w images were acquired with a repetition time/echo time (TR/TE) of 12 s/156 ms, SENSE 2.11/2.58 (axial/sagittal) with an in-plane resolution of 0.8 × 0.8 mm, slice thickness of 1.6 mm and overlap of 0.8 mm. Images were motion-corrected and super-resolved to produce a final resolution of 0.5 mm isotropic. Diffusion images were acquired over a spherically optimized set of directions on three shells (b = 400, 1000, and 2600 s/mm^2^). A total of 300 volumes were acquired per subject, including 20 with b = 0 s/mm^2^, across four acquisition subsets (two pairs with opposing phase encoding polarities). For each volume, 64 interleaved overlapping slices were acquired (in-plane resolution = 1.5 mm, thickness = 3 mm, and overlap = 1.5 mm). The data were then super-resolved along the slice direction to achieve an isotropic resolution of 1.5 mm^3^ and preprocessed to correct for motion and distortions ^[66, 67]^.

The dHCP data release includes an assessment of incidental findings scored 1 to 5, i.e., the "radiology score," with larger values indicating larger or more clinically significant incidental findings. Here, we used this scoring system to exclude subjects with a score greater than 3 (3 indicates "incidental findings with unlikely clinical significance but possible analysis significance"). The subjects with more than 10% of scan volumes excessive head movement or whose images were acquired after four PMW were also excluded. Only full-term neonates were included in the further analysis. Finally, 242 term neonates were included in this study (138 males; mean scan age, 40.30 PMW; scan age range, 37.14 -43.57 PMW).

#### Image processing

The pipeline used for processing the structural data was fully described in a previous publication ^[68]^. In summary, motion-corrected, reconstructed T2w images were first bias-corrected and brain-extracted. Following this, images were segmented into different tissue types using the Draw-EM algorithm. Next, topologically correct white matter surfaces were fit first to the grey-white tissue boundary and then to the grey-white interface of the MR intensities. Pial surfaces were generated by expanding each white matter mesh toward the grey-cerebrospinal fluid interface. Separately, an inflated surface was generated from the white surface through a process of inflation and smoothing. This inflated surface was then projected onto a sphere for surface registration. The surfaces and cortical metrics from each neonate were further nonlinearly aligned to the dHCP symmetric template (40 PMW) ^[69]^ using multimodal surface matching (MSM) ^[70]^.

The preprocessing of the diffusion data included denoising ^[71]^, correction for Gibbs ringing ^[72]^, distortion correction ^[73]^, SHARD motion correction ^[74]^, correction for intensity inhomogeneities ^[75]^, and linearly alignment to the T2-weighted space. The concatenation of the diffusion-to-T2 and T2-to-40-week-template transforms allowed transformation between diffusion and template space. DTIFIT was then used to fit a diffusion tensor model. The probability distributions of the fiber orientation distribution were estimated using Bedpostx ^[76]^.

#### Participants selection

The infants whose birth and scan age differed by less than seven days were used in the further analysis (*n* = 242, 138 males). The infants who had two scans and the second scan age was older than 40 PMW were used in the longitudinal analysis (*n* = 36, 20 males; the birth age ranged from 25.42 to 36.57 PMW; the first scan age ranged from 28.29 to 37.43 PMW; the second scan age ranged from 40.00 to 44.86 PMW).

### Connectopy mapping

#### Tractography

To map the whole-brain connectivity pattern, we performed probabilistic tractography using FSL’s probtrackx2 accelerated by using GPUs ^[77, 78]^. Specifically, the white surface was set as a seed region tracking to the rest of the brain with the ventricles removed. The pial surface was used as a stop mask to prevent streamlines from crossing sulci. Each vertex was sampled 5000 times (5000 trackings) based on the orientation probability model for each voxel, with a curvature threshold of 0.2, a step length of 0.5 mm, and a number of steps of 3200. This resulted in a (*whole-surface vertices*) × (*whole-brain voxels*) matrix for further analysis. We checked the preprocessing images and tractography results of each subject to ensure the accuracy of our analysis. Specifically, the reconstructed surfaces were visually inspected, and the transforms between different spaces were checked carefully. Tractography results were also visually checked to see if any streamline crossed the sulci.

#### Tractogram covariance analysis

We calculated the tractogram covariance (TC) matrix to characterize the structural connectivity similarity profile. The vertex profiles underwent pairwise Pearson correlations, controlling for the average whole-cortex profile. For a given pair of vertices, *i* and *j*, TC was calculated as

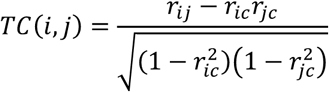

in which *r_ij_* was the Pearson correlation coefficient of the structural connectivity profile at vertices *i* and *j*, *r_ic_* was the correlation of the structural connectivity profile at vertex *i* with the average connectivity profile across the whole cortex, and *r_jc_* was the correlation of the structural connectivity profile at vertex *j* with the average connectivity profile across the entire cortex. A symmetric ∼30k×30k TC matrix was produced for each subject. Then, the TC matrices of all the subjects for each week were averaged separately for the left and right hemispheres to get a group-level TC matrix for each hemisphere. In line with a previous study ^[79]^, the TC matrix was proportionally thresholded at 90% per row, with elements above zero retained to remove negative connections.

#### Connectopy decomposition

Using a cosine affinity kernel, the TC matrix was transformed into a nonnegative square symmetric affinity matrix. Then, diffusion map embedding was implemented to identify the principal gradient components ^[28]^. Diffusion map embedding is a nonlinear manifold learning technique that maps cortical gradients ^[80]^. Along these gradients, cortical vertices with similar connectivity profiles are embedded along the axis. Diffusion map embedding is relatively robust to noise and less computationally expensive than other nonlinear manifold learning techniques.

Global connectopies were computed separately for the left and right hemispheres with two key parameters: *α* controls the influence of the density of sampling points on the manifold, and *t* controls the scale of the eigenvalues. Here, we set *α* = 0.5 and *t* = 0 as recommended ^[80]^. The first three global connectopies were chosen to map onto the cortical surface for further analysis. We also calculated two global measures, i.e., explanation ratio (i.e., the variance accounted for by the global connectopies) and gradient range (i.e., the difference between the values of the positive and negative extremes of the gradient axis). Each neonate’s global connectopies were calculated using the same methods.

### Contributions of thalamic projection tracts

#### Reconstruction of thalamic projection tracts

To investigate how the global connectopies are related to the underlying white matter tracts, we reconstructed thalamic projection tracts using a pre-trained deep-learning model, TractSeg ^[31]^. Specifically, we reconstructed nine thalamic projection tracts (anterior thalamic radiation: ATR; superior thalamic radiation: STR; thalamo-prefrontal tract: T PREF; thalamo-premotor tract: T PREM; thalamo-precentral tract: T PREC; thalamo-postcentral tract: T POSTC; thalamo-parietal tract: T PAR; thalamo-occipital tract: T OCC; optic radiation: OR).

#### Virtual lesion analysis

We assessed the importance of the thalamic projection tracts by performing a virtual lesion of the thalamocortical tracts. Specifically, we set the connectivity probability values of all the thalamocortical tracts (the same as above) in the tractogram to zero. Due to the ultra-high computational cost of vertex-wise analyses, we conducted the subsequent calculations at the ROI level. The new global connectopies were calculated using the same algorithm as the original ones.

## Data availability

The infant datasets are available at https://www.developingconnectome.org/. Data from BrainSpan can be downloaded at https://www.brainspan.org/.

## Code availability

This study utilized the following open software and code: dHCP structural pipeline: https://github.com/BioMedIA/dhcp-structural-pipeline, dHCP diffusion SHARD pipeline: https://github.com/dchristiaens/dhcp-shard-pipeline, FreeSurfer v6.0: http://surfer.nmr.mgh.harvard.edu/, FSL v6.0.7: https://fsl.fmrib.ox.ac.uk/fsl/fslwiki, MSM: https://github.com/ecr05/MSM_HOCR/releases, Connectome Workbench: https://www.humanconnectome.org/software/connectome-workbench, BrainSpace: https://brainspace.readthedocs.io/. The graphical representation of symbols was created with BioRender.com.

## Acknowledgments

This work was partially supported by STI2030-Major Projects (Grant No. 2021ZD0200203), the Natural Science Foundation of China (Grant Nos. 82072099, 82202253, 62250058), and the China Postdoctoral Science Foundation (2022M722915). We also thank the Developing Human Connectome Project for providing the publicly available neonatal brain MRI data for the validation analysis. Data were provided by the Developing Human Connectome Project, KCL-Imperial-Oxford Consortium, funded by the European Research Council under the European Union Seventh Framework Programme (FP/2007-2013)/ERC Grant Agreement no. [319456]. The authors appreciate the English language and editing assistance of Rhoda E. and Edmund F. Perozzi, PhDs.

## Supplementary Figures

**Figure S1.**
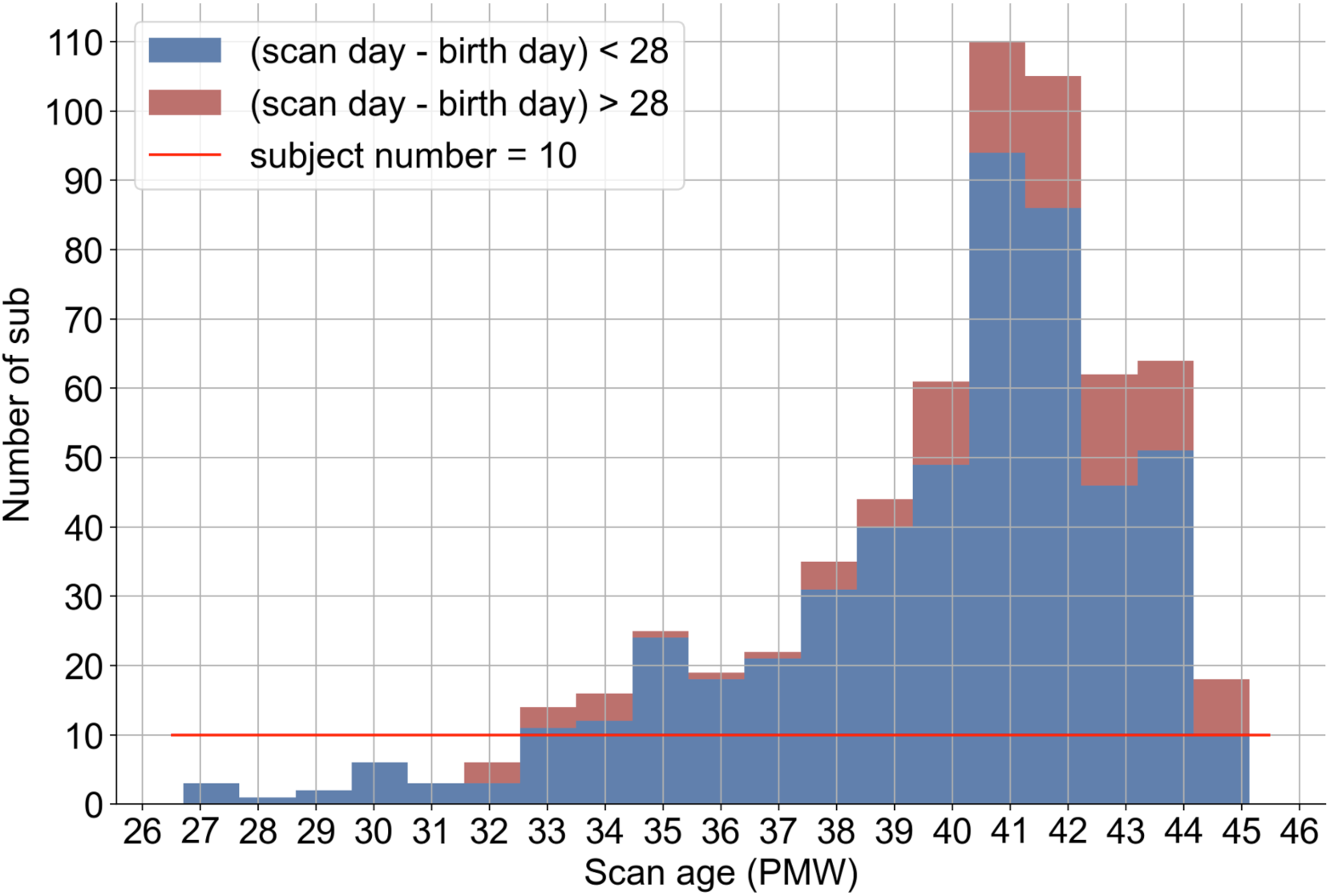
Study demographics and inclusion criteria for neonatal subjects. To ensure the robustness of our findings, our analysis was restricted to subjects with imaging data acquired within a four-week window and was further limited to postmenstrual age weeks with a participant count exceeding ten.

**Figure S2.**
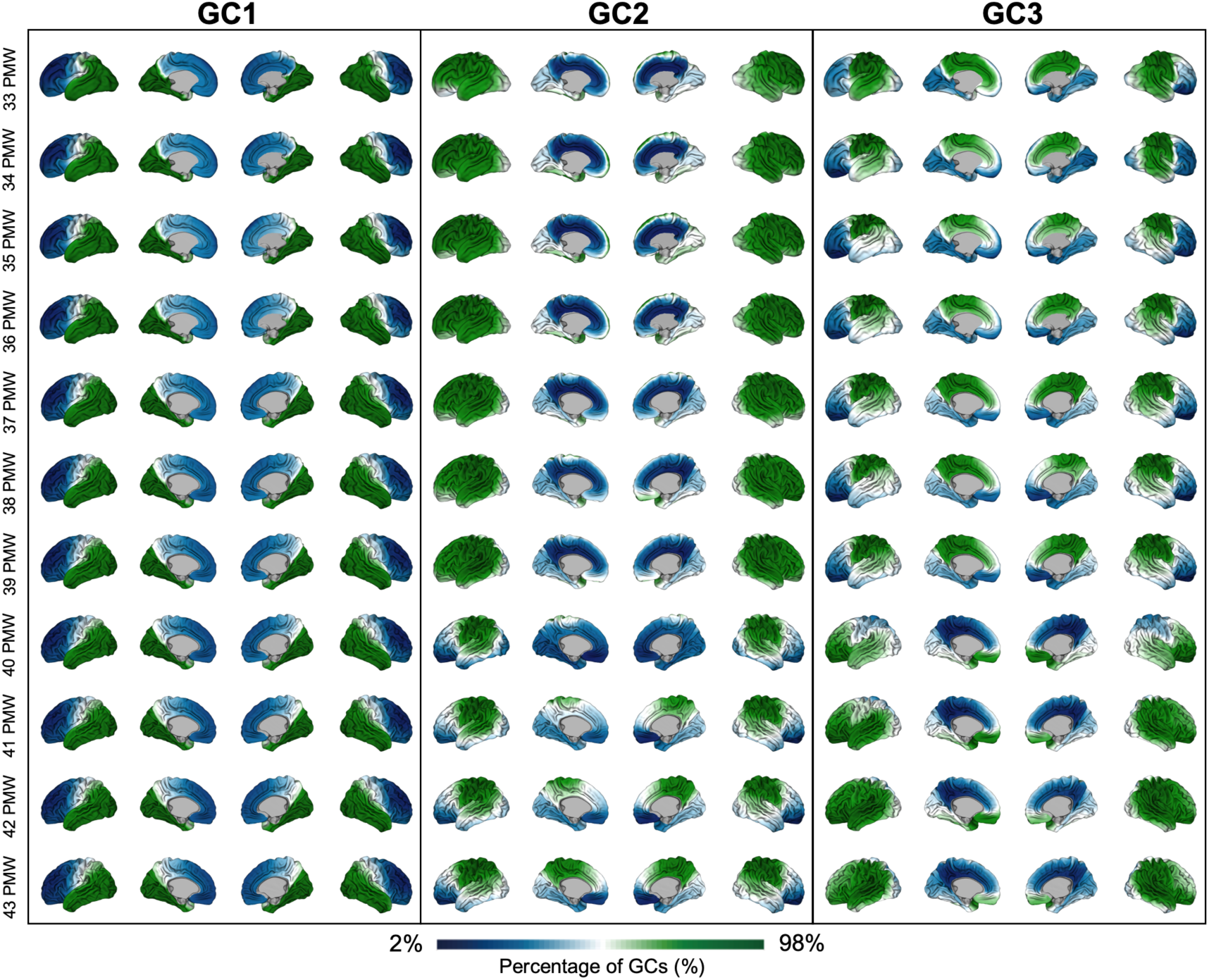
The three global connectopies of human infants from 33 to 43 PMW.

**Figure S3.**
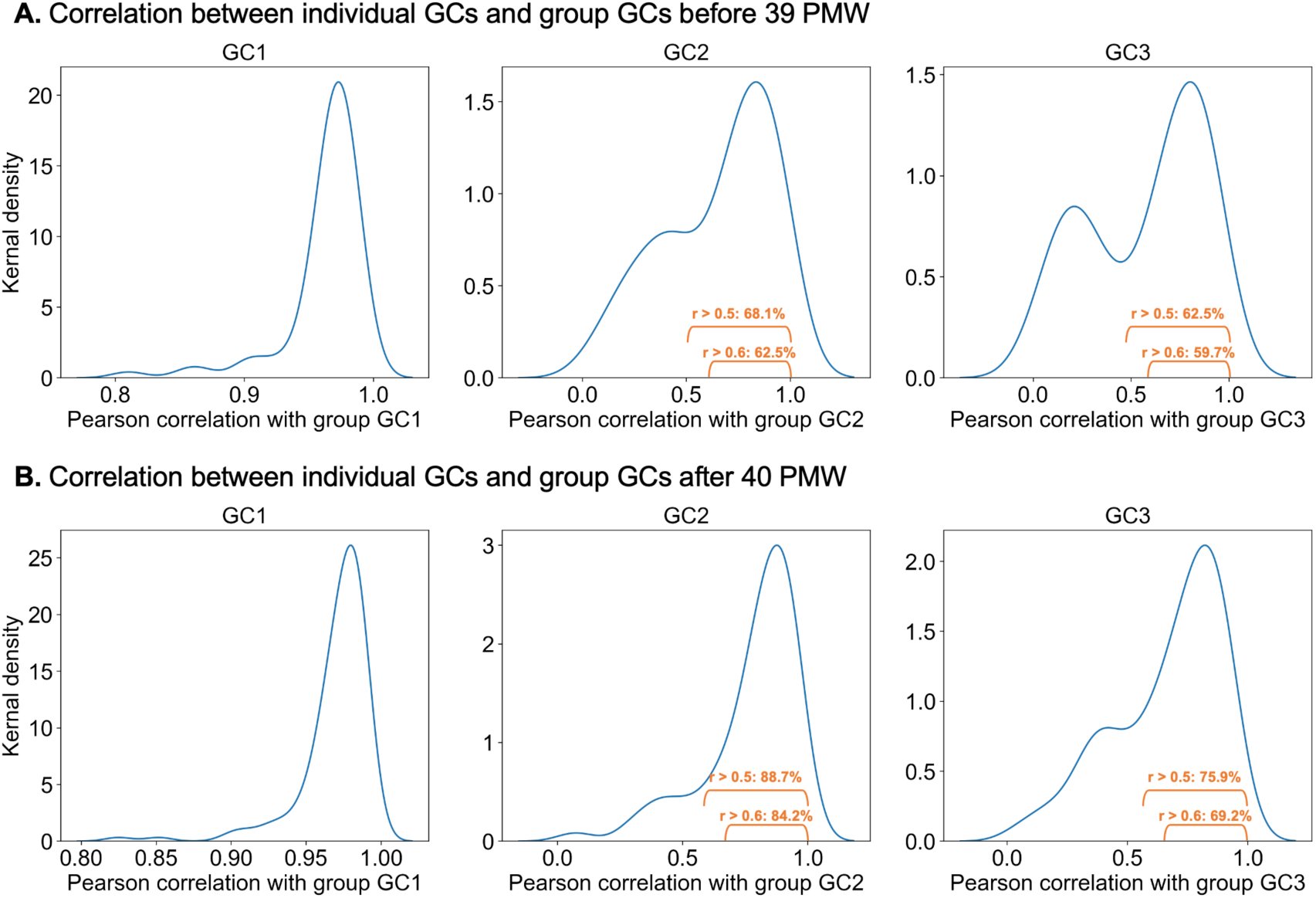
Individual global connectopies exhibited a high correlation with group-level global connectopies. **(A)** The distribution of Pearson correlation coefficients between each individual’s GCs under 39 PMW and the age-matched group GCs. **Left panel:** The GC1 of all infants is highly correlated at the group level (*r* > 0.8); **Middle panel:** More than two-thirds of infants show a correlation greater than 0.5 between their GC2 and the group-level GC2; **Right panel:** 62.5% of infants show a correlation greater than 0.5 between their GC2 and the group-level GC2. **(B)** The distribution of Pearson correlation coefficients between each individual’s GCs after 40 PMW and the age-matched group GCs. **Left panel:** The GC1 of all infants is highly correlated at the group level (*r* > 0.8); **Middle panel:** 88.7% of infants show a correlation greater than 0.5 between their GC2 and the group-level GC2; **Right panel:** 75.9% of infants show a correlation greater than 0.5 between their GC2 and the group-level GC2. The GC1 of all infants is highly correlated at the group level.

